# Toward defining loss functions in neuroscience: an XOR-based neuronal mechanism for local learning

**DOI:** 10.64898/2026.03.16.712061

**Authors:** María Peña Fernández, Lara Lloret Iglesias, Jesús Marco de Lucas

**Affiliations:** Advanced Computing and e-Science Group, Instituto de Física de Cantabria (IFCA) CSIC-Universidad de Cantabria, Santander, ES 39005 SPAIN

**Keywords:** local learning rule, credit assignment, biologically plausible learning, XOR motif, prediction error, autoencoder, feedback alignment

## Abstract

Of the three components often used to map deep learning onto neuroscience (architecture, learning rule and loss function), the biological counterpart of the loss is the least understood. For binary signals, the mismatch between an externally evoked response and its internally generated reconstruction is the XOR function, and its two asymmetric components, undershoot and overshoot, form a signed error that also indicates the direction of the required change. In a companion study, an eight-neuron, Dale-compliant signed-XOR motif, a six-neuron XOR comparator with two feedback neurons of opposite identity, was found in several connectomes and shown in spiking simulations to compute this signed error. Here we ask whether this signed error can serve as an effective learning signal. We use the logical function of one comparator per output unit as the only error signal of a binary autoencoder, with fixed random sign feedback to the hidden layer. Without backpropagation, a fully connected autoencoder trained for five epochs on binarized MNIST reaches 97.7% bit-wise and 94.8% balanced reconstruction accuracy (five seeds), and a convolutional variant with 533 parameters reaches 96.3% and 89.2%; without the sign of the error, learning collapses. Controls show that performance is carried by the comparator-driven decoder: a suitably initialised fixed random encoder reconstructs digits about as well or better, encoder learning improves digit readout mainly in narrow hidden layers, and a gradient-trained reference reaches 99.3% and 97.3% balanced accuracy. The signed error thus suffices for the synapses it reaches directly; credit assignment to earlier layers remains open.

## 1 Introduction

When we compare how artificial intelligence systems operate with biological systems, although both learn from experience, it is not easy to establish an equivalence. In artificial neural networks, learning is guided by a well-defined loss function that measures the error between the output of the model being trained and the desired target. Credit assignment, i.e. the adjustment of the network weights, relies on global algorithms such as backpropagation, which follows the gradient of that loss function (Rumelhart et al. 1986).

In biological neural networks, in contrast, it is not clear what the equivalent of a loss function is, how it could be computed, and—given that backpropagation does not seem biologically plausible (Lillicrap et al. 2020)—what guides credit assignment, i.e. the adjustment of connections between neurons, so that performance improves.

Addressing this question is a key objective of NeuroAI research (Richards et al. 2019). Both neuroscientists and machine learning researchers are looking for plausible local learning rules (i.e. rules that use only information available at each synapse, without integrating information from the whole system) that allow artificial neural networks to approach the efficiency of the brain (Wu et al. 2022, Gilra & Gerstner 2017, Murray 2019, Whittington & Bogacz 2019). Several local alternatives to backpropagation have been proposed: random feedback alignment (Lillicrap et al. 2016) and its direct variant (Nøkland 2016), predictive-coding networks that approximate backpropagation (Whittington & Bogacz 2017), dendritic error computation in cortical microcircuits (Sacramento et al. 2018), equilibrium propagation (Scellier & Bengio 2017), eligibility-trace rules for spiking networks (Bellec et al. 2020), and burst-dependent plasticity (Payeur et al. 2021). These approaches specify how an error is transported to each synapse, but they generally assume that the error itself—the difference between a target and a prediction—is available as a graded quantity. How a neural circuit would compute that difference is rarely made explicit. Experimentally, neurons that signal a mismatch between expected and actual sensory input have been described in mouse primary visual cortex (Keller et al. 2012), with distinct, molecularly identifiable populations for positive and negative prediction errors (Keller & Mrsic-Flogel 2018, O’Toole et al. 2023); circuit models show that inhibitory plasticity in a canonical interneuron circuit can learn to produce such prediction-error neurons (Hertäg & Sprekeler 2020). In addition, the dendrites of human layer 2/3 pyramidal neurons have been shown to implement an XOR-like input–output function (Gidon et al. 2020).

One perspective draws on *predictive learning* (Rao & Ballard 1999) *and memory engrams. In previous work (Marco 2023) we proposed that the brain might form compressed latent representations of sensory input (like the hidden layer of an autoencoder) and learn by matching the reconstruction to the original input, comparing external sensory signals with internal predictions. The key question is which neural process could implement* this comparison and signal an error that drives synaptic changes.

Such an error signal has two requirements. It must be local, readable by a synapse from the neurons it directly contacts, and it must carry a sign, telling the synapse whether to potentiate or to depress. We previously proposed that a six-neuron motif organised around a single inhibitory interneuron computes an exclusive OR (XOR) between two binary signals, for example an external input and an internal prediction of it (Peña et al. 2024) (Fig. 1a). Its output neuron fires when the two signals differ and stays silent when they match, so that matched activity is cancelled locally. The XOR output, however, signals only *whether* there is a mismatch, not its direction.

**Figure 1.**
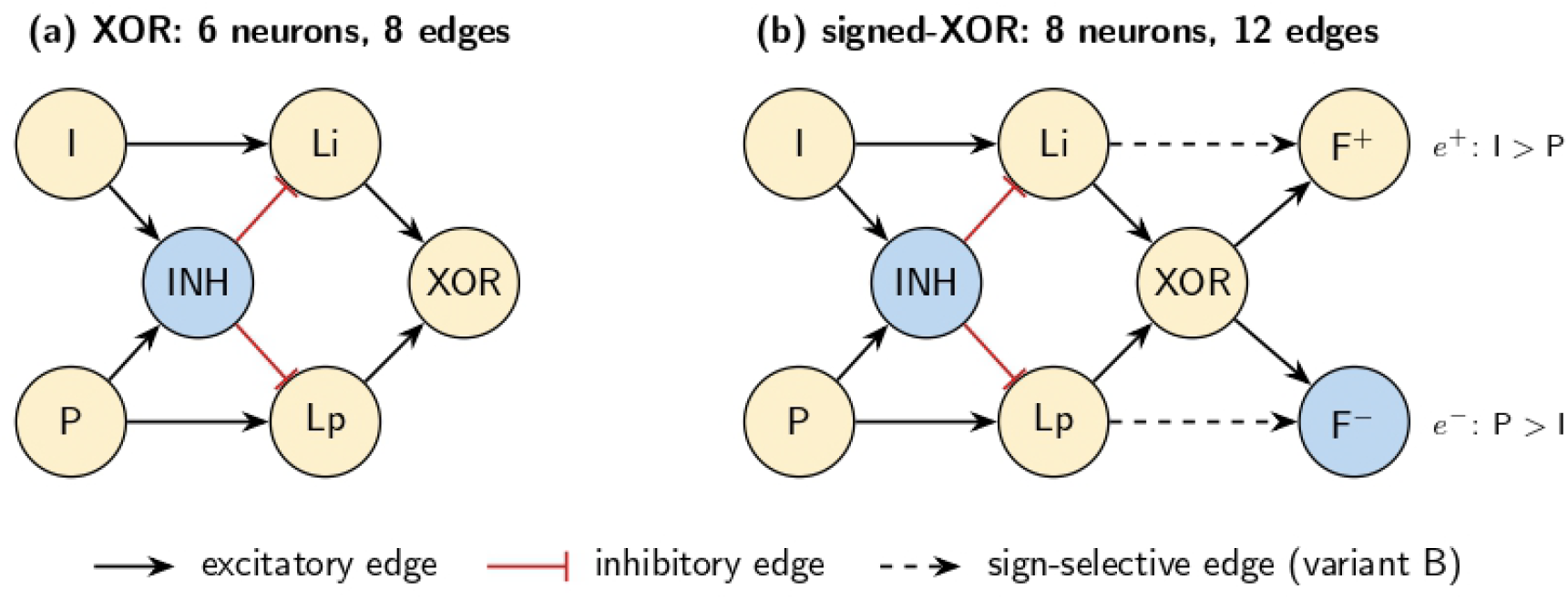
The XOR and signed-XOR motifs. (a) Six-neuron XOR motif (Peña et al. 2024): two excitatory inputs, I (e.g. external input) and P (e.g. internal prediction), two excitatory relays Li and Lp, an inhibitory neuron INH that receives both inputs and inhibits both relays, and the comparator XOR, which is active if and only if exactly one input is active. (b) Eight-neuron signed-XOR motif (Peña Fernández et al. 2026): two feedback neurons of opposite identity are added, F^+^ (excitatory), active when I exceeds P (undershoot, *e*^+^), and F^*−*^ (inhibitory), active when P exceeds I (overshoot, *e*^*−*^). Shown is the variant in which F^+^ and F^*−*^ read the relays (variant B of (Peña Fernández et al. 2026)); in variant A they read the inputs. Excitatory neurons in cream, inhibitory in blue. The autoencoders of this work use only the logical function of the motif (Section 2.2).

In a companion preprint (Peña Fernández et al. 2026), developed in parallel with the present work, we extended the XOR core with two feedback neurons of opposite identity: an excitatory neuron F^+^ that fires when the input exceeds the prediction and an inhibitory neuron F^*−*^ that fires when the prediction exceeds the input (the eight-neuron signed-XOR motif; Fig. 1b). That study formalises the motif, finds it in the connectomes of *C. elegans, Drosophila* and mouse visual cortex, where its enrichment is comparable to that of alternative wirings of the same neurons, and shows in spiking simulations of all 50 such wirings that only a small family computes the signed error, the signed-XOR among them. It thus establishes that a signed local error can be produced by a concrete, Dale-compliant microcircuit. It does not test whether the error it emits can train a network.

This is the question addressed here: *can the signed error produced by such a comparator serve as an effective learning signal in a larger neural network?* We answer it at the computational level. We do not simulate the spiking circuit; we take its function, the two components of the signed error, as given, and use one comparator per output unit as the only error signal of binary autoencoders trained with local update rules. Locally, this error is self-limiting: the comparator is active while its own bit is wrong and silent once it is right, a homeostatic condition in which no further change is triggered at that unit. Whether the changes triggered at individual units reduce the error of the whole network, and which synapses they can train, are empirical questions.

The contributions are: (i) a formulation of the per-unit XOR mismatch as a signed error whose two components correspond to the F^+^ and F^*−*^ channels of the signed-XOR motif, and which, for the synapses that the comparator reaches directly, supplies exactly the error of the classical perceptron rule (Section 2.2); (ii) a demonstration, over five random seeds, that this signal trains fully connected and convolutional binary autoencoders within five epochs, and that the sign of the error is necessary (Sections 3.1–3.2); and (iii) a set of control experiments that separates what the rule achieves from what the architecture provides: reconstruction is carried by the comparator-driven decoder, encoder learning with fixed random sign feedback improves the linear decodability of the trained digit classes mainly for the narrowest hidden layers, and the gap to a gradient-trained reference is quantified (Section 3.3). We therefore present the work as an evaluation of the capabilities and limits of a local neuronal error signal, not as a competitive optimisation technique.

## Materials and methods

We used the function of the signed-XOR comparator as the error signal of a simple neural network to test its ability to guide learning, analogous to how a loss function guides weight adjustments in machine learning. We implemented a one-hidden-layer autoencoder with binary neurons as a testbed. The network’s goal is to reconstruct its input at the output layer; each output unit is paired with one comparator, which compares the corresponding input bit with the activity of the output neuron and signals whether and in which direction they differ.

### 2.1 Network architectures

#### Fully connected autoencoder

The autoencoder consists of an input layer, one hidden layer (latent representation), and an output layer of the same size as the input (Fig. 2). For MNIST (LeCun et al. 1998), the input is a vector of 784 bits (28*×* 28binary pixels) and the hidden layer has 1024 neurons. Each neuron produces a binary output by thresholding its total input:

**Figure 2.**
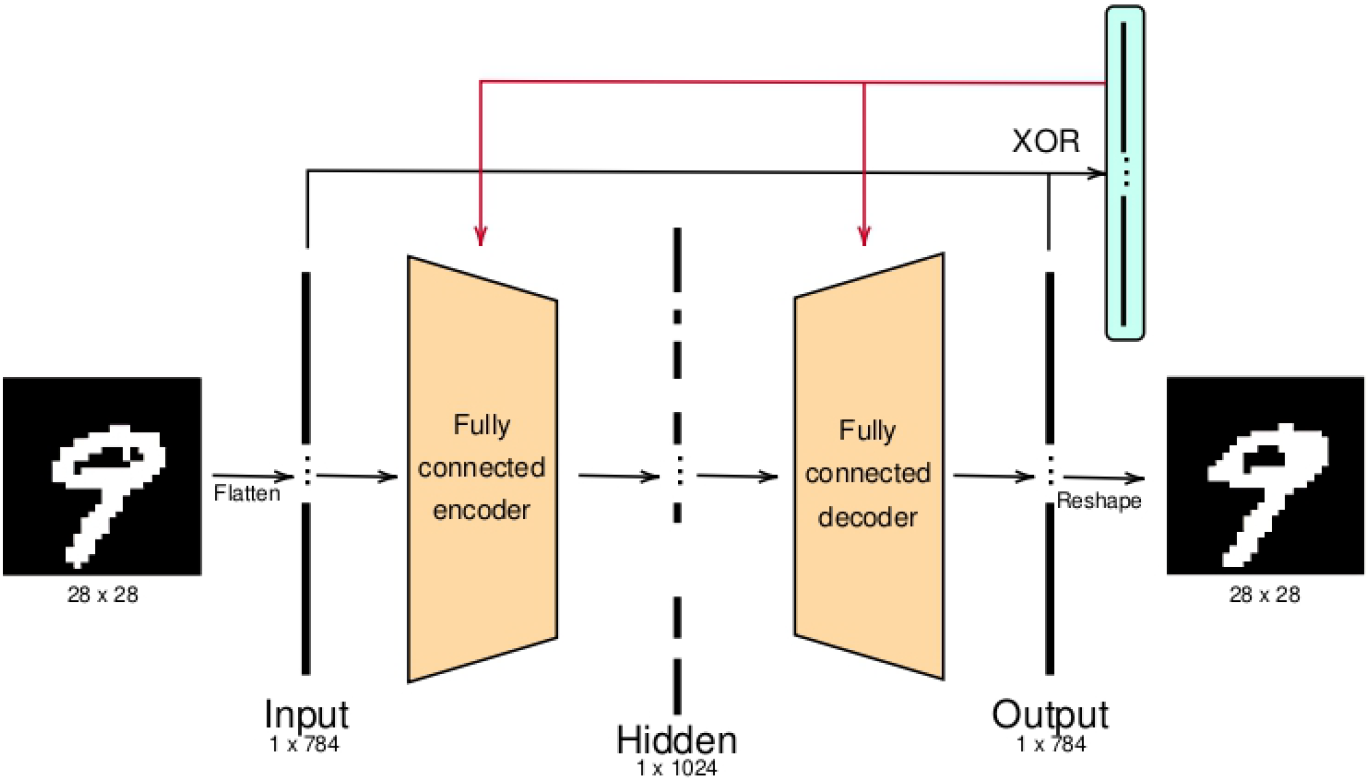
Architecture of the fully connected autoencoder. An array of XOR comparators, one per output unit, compares each input bit with its reconstruction.

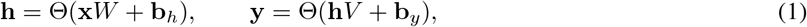

where **x***∈ {* 0, 1*}*^784^ is the input (row vector), *W ∈* ℝ^784*×*1024^ and *V ∈* ℝ^1024*×*784^ are the encoder and decoder weights, **b**_*h*_, **b**_*y*_ the biases, and Θ*u*=1if *u ≥* 0and 0otherwise. Weights are initialized from *N* 0, 0.01^2^and biases to zero. The network has 1 607 440 trainable parameters. No backpropagation or global error computation is used; learning is driven by local mismatch signals at the output.

#### Convolutional autoencoder

The encoder applies 28 filters of size 3*×* 3(stride 1, padding 1) followed by the threshold Θand 4*×* 4max-pooling, giving a latent code of 28*×* 7*×* 7=1372binary units, comparable in size to the fully connected network (Fig. 3). The decoder first restores each pooled unit to the position of the maximum within its 4*×*4window (max-unpooling with the pooling indices), then applies a single 3*×* 3filter and the threshold. The network has 533 trainable parameters (252 encoder weights and 28 biases; 252 decoder weights and one bias). The pooling indices carry information about where each maximum occurred that is not part of the 1372-unit code; we therefore also trained a variant in which unpooling is replaced by nearest-neighbour upsampling, which removes this side channel (Section 3.2).

**Figure 3.**
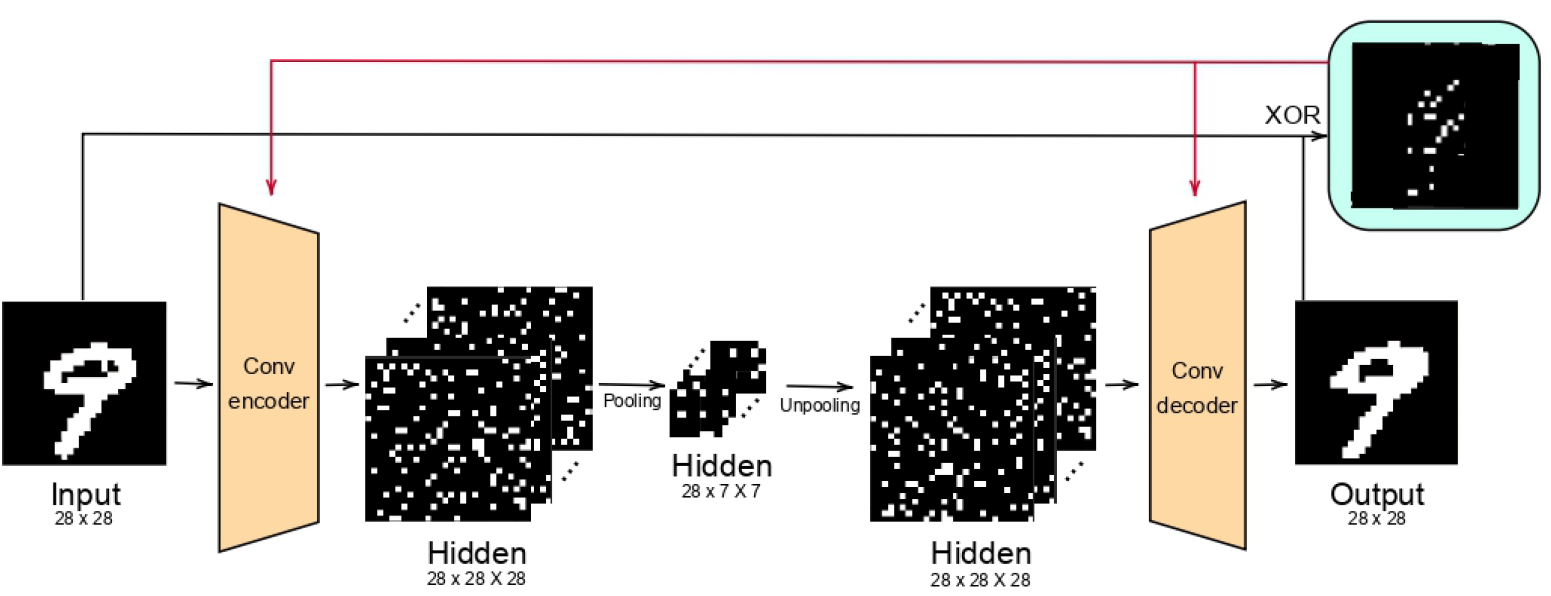
Architecture of the convolutional autoencoder.

### 2.2 Signed-XOR local learning rule

When an input pattern **x** is presented, the network produces a reconstruction **y** (Eq. 1). For each output unit *i* the comparator computes

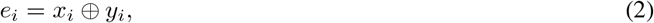

which is 1 if the reconstruction of bit *i* is wrong and 0 otherwise. The XOR output indicates *whether* a unit must change, but not in which direction. The direction is given by which of the two streams is active, i.e. by the two asymmetric components of XOR,

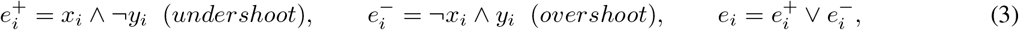

each of which can be computed by an excitatory input gated by an inhibitory one. We refer to their difference,

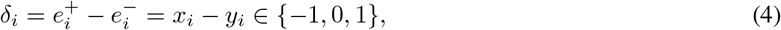

as the *signed XOR* error. Its magnitude is the XOR output and its sign identifies the active component. In the signed-XOR motif (Fig. 1b), 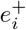 and 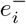 are the activities of the feedback neurons F^+^ and F^*−*^, so that the sign is delivered without violating Dale’s principle; the decoder update below, written in (Peña Fernández et al. 2026) as Δ*V*_*ij*_ =*η h*_*j*_ *e*_*i*_ 2*x*_*i*_ *−* 1, is the same rule.

#### Level of description

The autoencoders implement the logical function of the comparators, 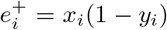 and 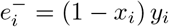, not their circuitry: no spiking motif is simulated, and the comparators have no dynamics, delays or failures. Dale’s principle applies to the motif that would produce the error; the weights *W, V* and *B* of the autoencoder take both signs and are not constrained by it. The connectivity of the motif and its spiking dynamics are the subject of (Peña Fernández et al. 2026); the present work addresses the third level, the use of the signed error as a learning signal in a network.

The decoder is updated with a Hebbian-like rule gated by the signed error: if *δ*_*i*_ =+1(the output under-shot the target), the connections from all active hidden neurons to output *i* are potentiated; if *δ*_*i*_ =*−*1(the output fired when it should not have), those connections are depressed; if *δ*_*i*_ =0, they are left unchanged. For a mini-batch of *N* patterns, with the rows of *X, H* and Δholding **x, h** and *δ*,

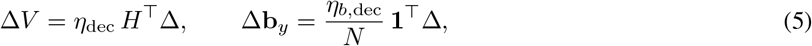

i.e. weight changes are summed over the mini-batch and bias changes are averaged. Each update uses only quantities available at the synapse: presynaptic activity *h*_*j*_ and the comparator output for the postsynaptic unit *i*. For a threshold output unit this is the perceptron (delta) rule (Rosenblatt 1962, Widrow & Hoff 1960), with the comparator supplying the error that the rule requires. The point of the present work is therefore not a new learning rule, but that a concrete neuronal circuit can supply the signals that this classical rule needs, and how far that local realisation goes when it is the only error signal of a network.

Updating the encoder weights *W* is harder because the error is detected at the output. To avoid transporting the decoder weights backwards, we use fixed random feedback, as in feedback alignment (Lillicrap et al. 2016): a random matrix *B ∈* ℝ^784*×*1024^ (the shape of *V* ^*⊤*^) with entries drawn once from *N* 0, 1*/*784and never trained projects the output error onto the hidden units. Since there is a single hidden layer, this coincides with direct feedback alignment (Nøkland 2016). We keep only the sign of the projected error,

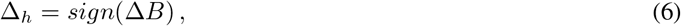

which serves as an estimate of the error direction at each hidden unit, and update

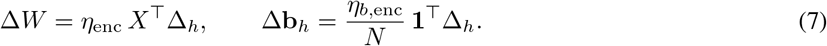

In the convolutional network the same rules are applied to the shared filter weights, summing over spatial positions and over the mini-batch and averaging bias changes over both; the feedback *sign (δ * B)*is obtained by convolving the output error with 28 fixed random 3*×*3kernels (entries from *N* 0, 1*/*9) and is applied at every pre-pooling position. The learning rates are *η*_dec_ =0.05, *η*_enc_ =0.02, *η*_*b*,dec_ =0.02and *η*_*b*,enc_ =0.01.

### 2.3 Training protocol and metrics

MNIST and EMNIST (balanced split, 47 classes, in the transposed orientation in which torchvision provides it, as in the original code; (Cohen et al. 2017)) were binarized with a threshold of 0.5 on intensities scaled to 0, 1. The autoencoders were trained on the 60 000 MNIST training images for five epochs in mini-batches of 256. Each mini-batch is presented repeatedly: after an update the network reconstructs the same mini-batch again and, while any bit is still wrong, applies a further update, up to three updates per mini-batch. In all runs three updates were applied to every mini-batch, giving 3525 weight updates in five epochs. These repeated presentations are part of training only; all reported test metrics come from a single forward pass of the trained network. No external reward or global error was provided at any point; learning is entirely unsupervised except for the linear readouts (Section 2.4).

We report *bit-wise reconstruction accuracy* (the fraction of input bits correctly reproduced) on the MNIST and EMNIST test sets; EMNIST (balanced split) contains both digits and letters and was never seen during training; in the orientation used here its images are, in addition, transposed with respect to MNIST. EMNIST results therefore measure transfer to a different distribution that combines new characters with a geometric transformation, not generalisation to new characters alone. Since the data are highly skewed (13.4% of MNIST and 17.0% of EMNIST test pixels are on, so an all-zero output already gives 86.6% and 83.0% bit-wise accuracy), we also report *balanced accuracy* (the mean of the accuracies on active and inactive pixels) and the F1-score for active pixels. Finally, we report the Learned Perceptual Image Patch Similarity (LPIPS) (Zhang et al. 2018) with the VGG backbone, computed on the binary images replicated to three channels and averaged over all test images (lower is better).

Every configuration was trained with five random seeds, which set the initial weights, the feedback matrix and the order of the mini-batches; we report mean *±* standard deviation over seeds. Configurations of the same architecture share the initialisation and the batch order for a given seed, so comparisons between them are also made seed by seed (paired differences). The control experiments use a re-implementation of the training code that is bit-identical to the original one with default settings, which we verified directly (Data availability).

### 2.4 Linear readouts of the latent code

To evaluate the latent representation, we trained linear classifiers on the hidden-layer activations of the frozen encoder to predict the class label. The first is a multi-class perceptron trained with a local rule: the predicted class is the argmax of the linear outputs, and on each mini-batch of 256 the weights of the correct class are incremented and those of the wrongly predicted class decremented (*η* =0.05for weights, 0.02for biases; five passes through the training set). The second is a least-squares (ridge) classifier fitted to one-hot targets, with the regularisation *λ ∈ {* 10^*−*2^, …, 10^3^*}* selected on a random 10% of the training set and the classifier then refitted on the full training set; it estimates how much class information is linearly available in the code. For the reference encoders, their initial (random) encoders and the raw pixels, extending the *λ* grid to 10^6^ changes least-squares accuracies by at most 0.4 points (raw-pixel EMNIST: 52.2% to 52.5%); some other configurations also select the largest *λ* of the grid, so their least-squares accuracies may be slightly underestimated. For EMNIST, a 47-class readout is trained on the EMNIST training codes produced by the encoder trained on MNIST and evaluated on the EMNIST test set. Both readouts are also computed on the encoder before training (a fixed random encoder) and on the raw pixels.

### 2.5 Control experiments

To identify what the signed-XOR rule contributes, we trained the following variants, each with the same five seeds: (a) *fixed random encoder*: only the decoder learns; (b) *unsigned error*: *δ*_*i*_ replaced by *e*_*i*_ =|*x*_*i*_*− y*_*i*_| ; (c) *graded feedback*: *sign*(Δ*B*)replaced by Δ*B*, either as is or rescaled so that its mean absolute value is 1, the step size of the sign; (d) *no bias learning*; (e) *single update per mini-batch*, for five epochs and for fifteen epochs (the latter matches the 3525 weight updates of the reference); (f) *bottleneck sweep*: hidden layers of 64 to 2048 units, with trained and with fixed encoders; (g) *training on random images*: Bernoulli images with the pixel density of MNIST, which contain no digit structure; (h) for the convolutional network, a small negative initial encoder bias (*b*_*h*_ =*−*10^*−*3^; see Section 3.3) and nearest-neighbour upsampling instead of unpooling. As a gradient-based reference, the same architectures were trained by backpropagation through a straight-through estimator (STE) of the threshold (Bengio et al. 2013), which passes the gradient where |*u*| *≤* 1(Adam (Kingma & Ba 2015), binary cross-entropy on the pre-threshold output, five epochs). The STE makes one update per mini-batch (1175 in five epochs, a third of those of the signed-XOR rule). Its learning rate was selected by the final training loss, without using test data, from 10^*−*4^ to 10^*−*2^ for the fully connected network and from 10^*−*4^ to 3*×*10^*−*1^ for the convolutional one; the selected values (3*×*10^*−*3^ and 10^*−*1^) lie inside the explored ranges.

## 3 Results

### 3.1 Learning with local signed-XOR error signals

Using the fully connected autoencoder, we trained the network with the XOR comparators as the only source of error. The network learned to reconstruct its inputs (Table 1, Fig. 4): backpropagation of errors is not required for this task, and a collection of local comparators, one per output unit, complemented by fixed random feedback to the hidden layer, suffices.

**Table 1.**
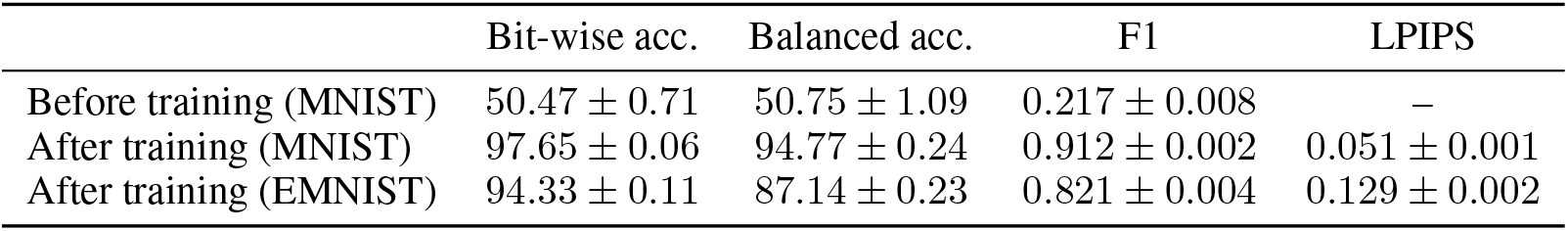
Reconstruction metrics of the fully connected autoencoder before and after five epochs of training on MNIST (mean *±* s.d. over five seeds). Accuracies in %. Higher is better for accuracy and F1; lower is better for LPIPS. An all-zero output gives 86.6% (MNIST) and 83.0% (EMNIST) bit-wise accuracy and 50% balanced accuracy.

|  | Bit-wise acc. | Balanced acc. | F1 | LPIPS |
| --- | --- | --- | --- | --- |
| Before training (MNIST) | 50.47 ± 0.71 | 50.75 ± 1.09 | 0.217 ± 0.008 | – |
| After training (MNIST) | 97.65 ± 0.06 | 94.77 ± 0.24 | 0.912 ± 0.002 | 0.051 ± 0.001 |
| After training (EMNIST) | 94.33 ± 0.11 | 87.14 ± 0.23 | 0.821 ± 0.004 | 0.129 ± 0.002 |

**Figure 4.**
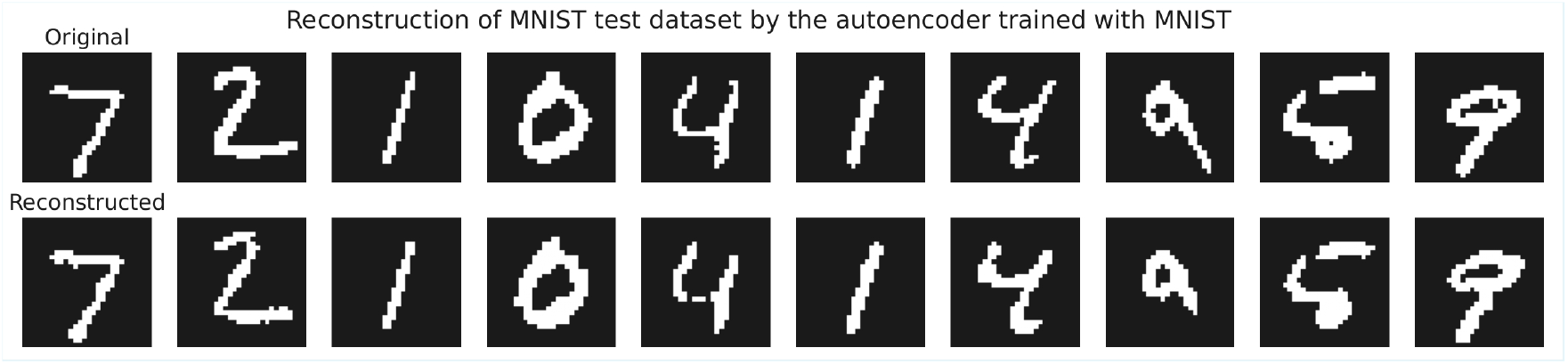
Reconstructions of MNIST test images after five training epochs of the fully connected autoencoder (one training run).

### 3.2 Reconstruction performance

Over the course of unsupervised training on MNIST, the reconstruction became progressively closer to the input. The network did not settle on the trivial all-zero output, even though no strategy to address class imbalance was used: after five epochs it reached 97.65*±* 0.06%bit-wise and 94.77*±* 0.24%balanced accuracy, with F1 =0.912on active pixels. Similarly, in biological systems, neural circuits learn to represent rare signals despite sparse firing. Reconstruction is good but not exact: on average 18.4*±* 0.5of the 784 pixels of a test image are wrong, and only 0.35% of test images are reproduced without any error. On EMNIST the network reaches 94.33%bit-wise and 87.14%balanced accuracy; Section 3.3 shows that this transfer does not require the network to have learned the structure of digits.

#### 3.1.2 The sign of the error is necessary

Replacing the signed error *δ*_*i*_ by the unsigned XOR output *e*_*i*_ removes the information about the direction of change: every wrong output is then pushed towards 1. As expected, the network collapses to an almost all-ones output (bit-wise accuracy 13.6*±* 0.1%, balanced accuracy 50.1*±* 0.1%, recall of active pixels 1.000, precision 0.13, i.e. the fraction of active pixels), and the convolutional network to an exactly all-ones output (13.4%and 50.0%). The collapse follows from the form of the update, so it should be read as a demonstration that the two asymmetric components of XOR must be kept separate, not as an empirical surprise. Other aspects of the rule matter much less. Using the graded projection Δ*B* instead of its sign lowers balanced accuracy slightly, both as is (94.25*±* 0.27%) and rescaled to the step size of the sign (94.42*±* 0.15%; paired difference with the sign *−*0.35*±* 0.10points, lower in all five seeds). Disabling bias learning leaves the fully connected network unchanged (94.75%balanced accuracy) but makes the convolutional one unstable (bit-wise accuracy 76.9*±* 9.3%).

#### 3.1.3 Repeated updates per mini-batch

Allowing up to three updates per mini-batch (Section 2.3) increases test accuracy compared with a single update (97.65vs. 96.90*±* 0.10%bit-wise). This effect is entirely explained by the larger number of weight updates: a single update per mini-batch for fifteen epochs, which matches the 3525 updates of the reference, gives 97.63*±* 0.06%. The repeated presentations are therefore additional plasticity steps during training, not an iterative refinement of the output at inference time. In the convolutional network neither the number of updates per mini-batch nor the number of epochs changes the result beyond the variation between seeds (96.10*±* 0.43%with one update per mini-batch, 95.97*±* 0.65%with one update for fifteen epochs, 96.32*±* 0.40%for the reference).

### 3.2 Convolutional autoencoder

The XOR-based rule also trains the convolutional architecture within five epochs (Table 2, Fig. 5). Reconstructions are less accurate than with the fully connected network, particularly for white pixels (recall 0.79vs. 0.91), consistent with the imbalance between black and white pixels. This lower accuracy must be weighed against model size: 533 parameters instead of about 1.6 million. Before training, the accuracy of the convolutional network depends strongly on the initialisation (47.1*±* 17.4%bit-wise), for the reason discussed in Section 3.3.

**Figure 5.**
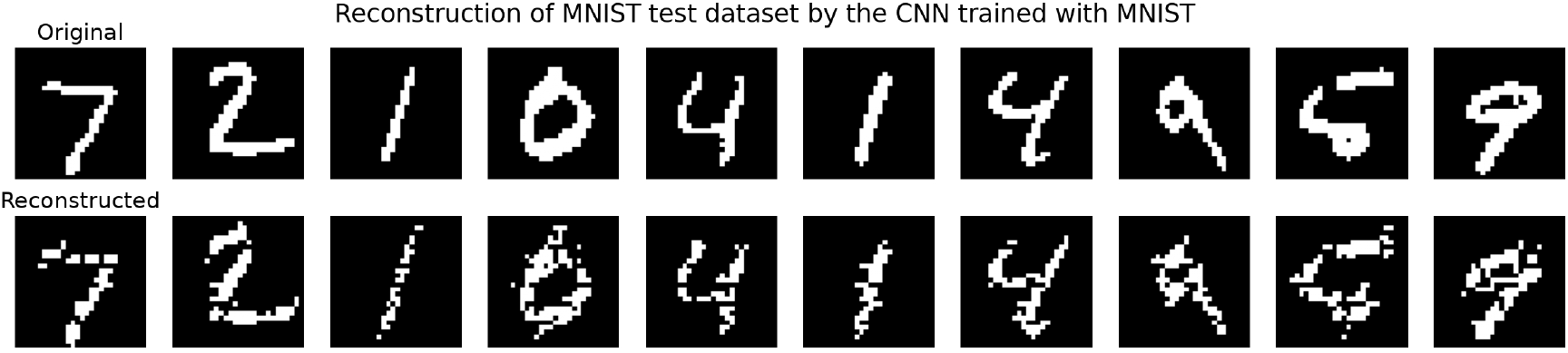
Reconstructions of MNIST test images after five training epochs of the convolutional autoencoder (one training run).

**Table 2.** Reconstruction metrics of the convolutional autoencoder before and after five epochs of training on MNIST, with max-unpooling (pooling indices passed to the decoder) and with nearest-neighbour upsampling (no indices). Same conventions as Table 1.

|  | Bit-wise acc. | Balanced acc. | F1 | LPIPS |
| --- | --- | --- | --- | --- |
| *Max-unpooling* |  |  |  |  |
| Before training (MNIST) | 47.10 ± 17.39 | 51.68 ± 8.04 | 0.235 ± 0.044 | – |
| After training (MNIST) | 96.32 ± 0.40 | 89.18 ± 0.97 | 0.853 ± 0.016 | 0.102 ± 0.010 |
| After training (EMNIST) | 95.34 ± 0.48 | 88.64 ± 0.95 | 0.851 ± 0.016 | 0.131 ± 0.011 |
| *Nearest-neighbour upsampling* |  |  |  |  |
| After training (MNIST) | 93.75 ± 0.38 | 86.69 ± 1.04 | 0.768 ± 0.015 | – |
| After training (EMNIST) | 93.02 ± 0.39 | 87.40 ± 0.83 | 0.794 ± 0.012 | – |

The MNIST–EMNIST gap in reconstruction is smaller for the convolutional network than for the fully connected one (1.0 vs. 3.3 points in bit-wise accuracy; 0.5 vs. 7.6 points in balanced accuracy). Part of the convolutional decoder’s information, however, does not pass through the latent code: the pooling indices tell it where each maximum occurred. On random images, the network with unpooling reconstructs 85.4*±* 4.0%of pixels in balanced accuracy, whereas with nearest-neighbour upsampling it reaches only 57.6*±* 0.9%(the fully connected network: 60.1%). Without the indices, MNIST reconstruction drops by 2.5 points in balanced accuracy (Table 2), but the MNIST–EMNIST gap remains small (EMNIST balanced accuracy is even slightly higher). The small gap is therefore not due to the index side channel; it is consistent with the locality of the filters, which do not depend on the identity of the characters.

### 3.3 What does the rule learn? Control experiments

Table 3 summarises the controls of Section 2.5.

**Table 3.**
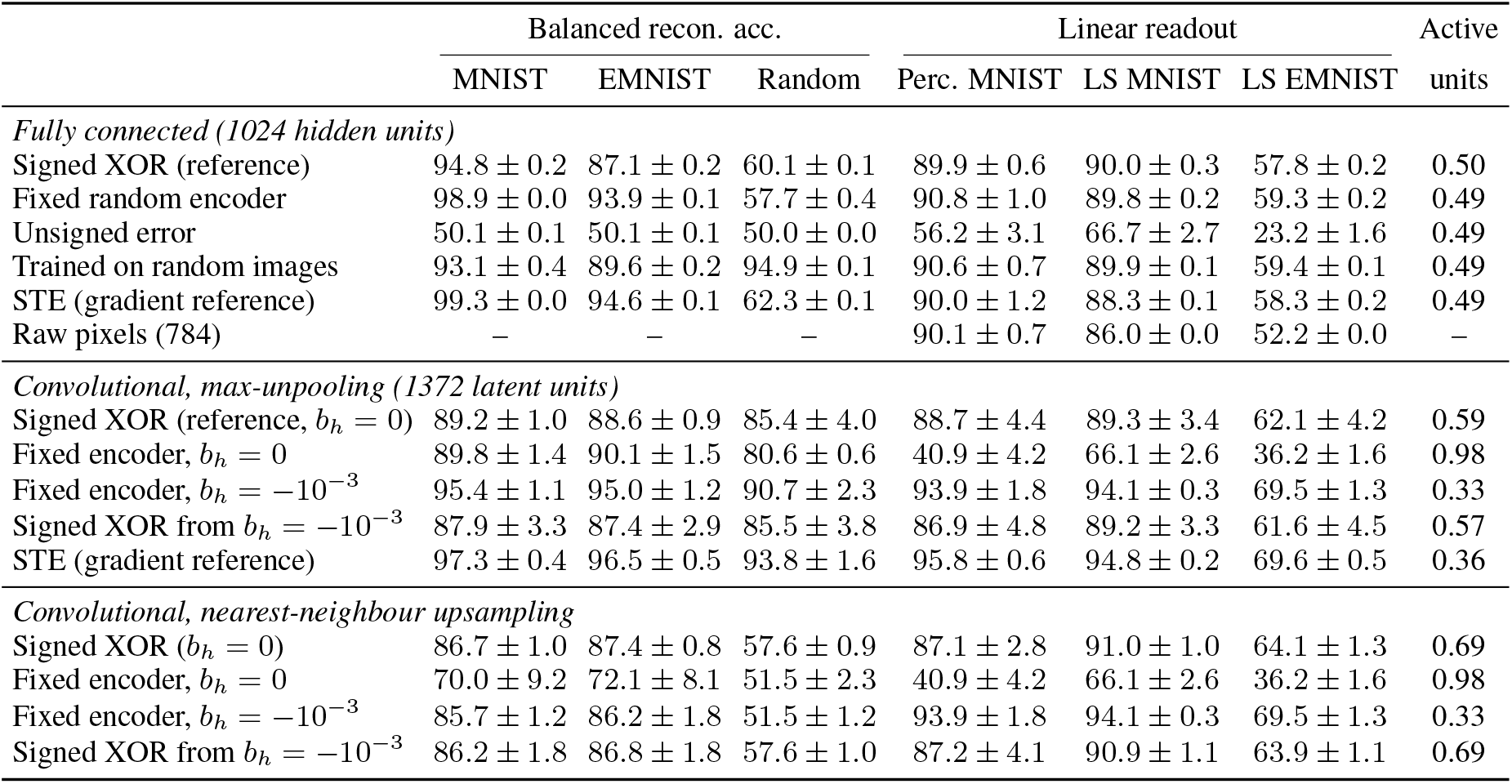
Control experiments (mean *±* s.d. over five seeds; accuracies in %). Balanced reconstruction accuracy on the MNIST, EMNIST and random-image test sets; linear readout of the latent code with the perceptron rule (Perc.) and with least squares (LS); fraction of active latent units. Fixed encoders keep their random initial weights. Rows that share an encoder (the fixed convolutional encoders with unpooling and with upsampling) share the readout columns.

|  | Balanced recon. acc. |  |  | Linear readout |  |  | Active units |
| --- | --- | --- | --- | --- | --- | --- | --- |
|  | MNIST | EMNIST | Random | Perc. MNIST | LS MNIST | LS EMNIST |  |
| *Fully connected (1024 hidden units)* |  |  |  |  |  |  |  |
| Signed XOR (reference) | 94.8 ± 0.2 | 87.1 ± 0.2 | 60.1 ± 0.1 | 89.9 ± 0.6 | 90.0 ± 0.3 | 57.8 ± 0.2 | 0.50 |
| Fixed random encoder | 98.9 ± 0.0 | 93.9 ± 0.1 | 57.7 ± 0.4 | 90.8 ± 1.0 | 89.8 ± 0.2 | 59.3 ± 0.2 | 0.49 |
| Unsigned error | 50.1 ± 0.1 | 50.1 ± 0.1 | 50.0 ± 0.0 | 56.2 ± 3.1 | 66.7 ± 2.7 | 23.2 ± 1.6 | 0.49 |
| Trained on random images | 93.1 ± 0.4 | 89.6 ± 0.2 | 94.9 ± 0.1 | 90.6 ± 0.7 | 89.9 ± 0.1 | 59.4 ± 0.1 | 0.49 |
| STE (gradient reference) | 99.3 ± 0.0 | 94.6 ± 0.1 | 62.3 ± 0.1 | 90.0 ± 1.2 | 88.3 ± 0.1 | 58.3 ± 0.2 | 0.49 |
| Raw pixels (784) | – | – | – | 90.1 ± 0.7 | 86.0 ± 0.0 | 52.2 ± 0.0 | – |
| *Convolutional, max-unpooling (1372 latent units)* |  |  |  |  |  |  |  |
| Signed XOR (reference, $b_h = 0$) | 89.2 ± 1.0 | 88.6 ± 0.9 | 85.4 ± 4.0 | 88.7 ± 4.4 | 89.3 ± 3.4 | 62.1 ± 4.2 | 0.59 |
| Fixed encoder, $b_h = 0$ | 89.8 ± 1.4 | 90.1 ± 1.5 | 80.6 ± 0.6 | 40.9 ± 4.2 | 66.1 ± 2.6 | 36.2 ± 1.6 | 0.98 |
| Fixed encoder, $b_h = -10^{-3}$ | 95.4 ± 1.1 | 95.0 ± 1.2 | 90.7 ± 2.3 | 93.9 ± 1.8 | 94.1 ± 0.3 | 69.5 ± 1.3 | 0.33 |
| Signed XOR from $b_h = -10^{-3}$ | 87.9 ± 3.3 | 87.4 ± 2.9 | 85.5 ± 3.8 | 86.9 ± 4.8 | 89.2 ± 3.3 | 61.6 ± 4.5 | 0.57 |
| STE (gradient reference) | 97.3 ± 0.4 | 96.5 ± 0.5 | 93.8 ± 1.6 | 95.8 ± 0.6 | 94.8 ± 0.2 | 69.6 ± 0.5 | 0.36 |
| *Convolutional, nearest-neighbour upsampling* |  |  |  |  |  |  |  |
| Signed XOR ($b_h = 0$) | 86.7 ± 1.0 | 87.4 ± 0.8 | 57.6 ± 0.9 | 87.1 ± 2.8 | 91.0 ± 1.0 | 64.1 ± 1.3 | 0.69 |
| Fixed encoder, $b_h = 0$ | 70.0 ± 9.2 | 72.1 ± 8.1 | 51.5 ± 2.3 | 40.9 ± 4.2 | 66.1 ± 2.6 | 36.2 ± 1.6 | 0.98 |
| Fixed encoder, $b_h = -10^{-3}$ | 85.7 ± 1.2 | 86.2 ± 1.8 | 51.5 ± 1.2 | 93.9 ± 1.8 | 94.1 ± 0.3 | 69.5 ± 1.3 | 0.33 |
| Signed XOR from $b_h = -10^{-3}$ | 86.2 ± 1.8 | 86.8 ± 1.8 | 57.6 ± 1.0 | 87.2 ± 4.1 | 90.9 ± 1.1 | 63.9 ± 1.1 | 0.69 |

#### Reconstruction is carried by the decoder

In the fully connected network, keeping the encoder fixed at its random initial weights and training only the decoder gives better reconstructions than training both: 98.95*±* 0.02%vs. 94.77%balanced accuracy on MNIST and 93.89%vs. 87.14%on EMNIST (better in all five seeds). On MNIST the advantage of the fixed encoder holds for all hidden-layer sizes from 128 to 2048 units (in all seeds); only with 64 units does the trained encoder reconstruct slightly better (84.10*±* 0.33%vs. 83.40*±* 0.68%; Fig. 6). On random images, in contrast, the trained encoder is better at every size, by 2 to 6 points, although it remains far from a network trained on random images (see below). The decoder rule is therefore sufficient for reconstructing digits, and encoder learning with random sign feedback does not help it for hidden layers of 128 units or more.

**Figure 6.**
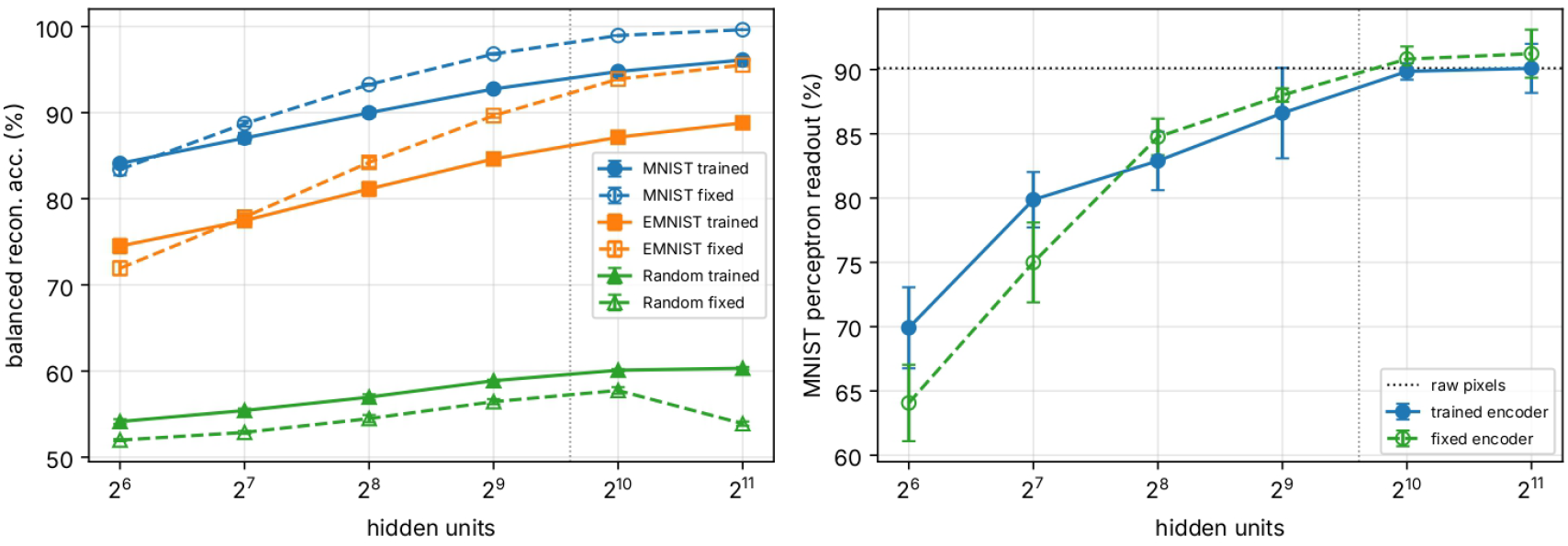
Fully connected autoencoder with hidden layers of 64 to 2048 units, with the encoder trained by the signed-XOR rule (solid) or kept at its random initial weights (dashed). Left: balanced reconstruction accuracy on MNIST, EMNIST and random images. Right: MNIST perceptron readout of the latent code; the dotted line is the readout of the raw pixels. The vertical line marks the input size (784). Mean *±* s.d. over five seeds.

#### Transfer to EMNIST does not require learned structure

A network trained on random images, which contain no digit structure, reconstructs MNIST with 93.11*±* 0.36%and EMNIST with 89.64*±*0.23%balanced accuracy, the latter above the network trained on MNIST (87.14%). With a hidden layer larger than the input, the decoder learns an approximately generic inverse of the encoder. This holds for the fully connected network; the convolutional network trained on random images reconstructs digits less well (80.47*±*3.04%on MNIST and 79.55*±* 3.04%on EMNIST). Training does specialise the network to its data: the network trained on MNIST reconstructs random images with only 60.10*±* 0.13%balanced accuracy, against 94.94*±* 0.13%for the network trained on random images.

#### Linear readout

A perceptron readout of the trained fully connected code classifies MNIST digits with 89.85*±* 0.65%accuracy (chance: 10%). The same readout of the code produced by the encoder before training gives 90.82*±* 0.98%, and of the raw pixels 90.10*±* 0.69%. The least-squares readouts of the trained and the random code are within one point of each other (90.05%and 89.78%); raw pixels are less separable by least squares (86.0%). On EMNIST (47 classes, chance *≈* 2.1%) the trained code is less decodable than the random one (least squares: 57.76*±*0.23%vs. 59.33*±* 0.20%; perceptron: 49.17*±* 2.38%vs. 58.19*±* 1.55%). With 1024 hidden units, therefore, the linear decodability of the code is not a product of learning: it is already available in the raw pixels and in a random projection of them. Linear decodability, and by extension the clustering of the hidden activations by digit class in a t-SNE projection (van der Maaten & Hinton 2008) that we reported in an earlier version of this work, cannot be taken as evidence of feature learning in this network. The per-class accuracy of the perceptron readout also varies with the seed: the worst class ranges from 74.4% to 85.7% across seeds.

#### Encoder learning helps digit readout mainly in narrow layers

The picture changes for the narrowest hidden layers (Fig. 6). With 64 hidden units the trained code gives a perceptron readout of 69.9*±*3.2%against 64.1*±*3.0%for the fixed random code, and a least-squares readout of 71.2*±* 1.2%against 66.6*±* 1.5%; with 128 units, 79.9*±* 2.2%against 75.0*±* 3.1%(perceptron) and 78.5*±* 0.7%against 75.7*±*0.2%(least squares, higher in all five seeds). From 256 units upward the perceptron readout favours the fixed code on average, and the least-squares advantage of the trained code falls below one point and vanishes at 2048 units (92.3%in both cases). The gain is also specific to the classes seen during training: on EMNIST, the trained code is less linearly decodable than the fixed one for every size from 128 units upward (least squares, lower in all seeds). Encoder learning with random sign feedback thus adapts the code to the training digits when the hidden layer is narrow, but it does not produce features that transfer to EMNIST.

#### Convolutional encoder and initialisation

In the convolutional network, training the encoder appears at first to be essential for the readout: the trained code gives 88.73*±*4.40%(perceptron) and the fixed random code only 40.87*±*4.19%. The reason is a degeneracy of the initialisation. With Θ0=1and zero biases, every filter is active on blank patches, so 98.5% of the units of the random code are on and the code is nearly saturated. A small negative bias (*b*_*h*_ =*−*10^*−*3^) silences blank patches: the fixed random encoder then has 33% active units, gives 95.38*±*1.13%balanced reconstruction accuracy and a readout of 93.93*±*1.81%(least squares 94.06%; EMNIST 69.50%), better on every measure than the trained encoder. Training the encoder from this initialisation lowers reconstruction to 87.95*±* 3.34%(lower in all five seeds) and the readout to 86.94*±* 4.75%. The benefit of encoder training in the reference convolutional network therefore comes from escaping the saturated initial code, not from learning features that a well-initialised random encoder lacks. Without the index side channel, the trained encoder reconstructs as well as the well-initialised fixed one (86.15vs. 85.72%; paired difference +0.43*±* 1.34points), but its code is again less decodable (least squares 90.86%vs. 94.06%, lower in all seeds).

#### Comparison with gradient training

Backpropagation through a straight-through estimator, with its learning rate tuned on the training loss, reconstructs better than the signed-XOR rule in both architectures: 99.34*±* 0.01%balanced accuracy on MNIST for the fully connected network (94.62%on EMNIST) and 97.32*±* 0.43%for the convolutional one (96.47%on EMNIST), against 94.77%and 89.18%. In the fully connected network the gap is closed to a large extent by the fixed random encoder (98.95%), in the convolutional network by the well-initialised fixed encoder (95.38%). Gradient training does not make the code markedly more linearly decodable either: the least-squares readout of the STE codes is close to that of the fixed random encoders (88.3%vs. 89.8%fully connected; 94.8%vs. 94.1%convolutional with *b*_*h*_ =*−*10^*−*3^). Linear decodability is therefore a weak probe of feature learning in this setting, whatever the learning rule.

## 4 Discussion

### 4.1 A neuronal comparator as the source of a local error

The connectomic and spiking study of the signed-XOR motif (Peña Fernández et al. 2026) identifies a candidate microcircuit that can generate a local error and deliver its sign through two channels of opposite identity. The present work tests the computational side of that hypothesis. Using only the function of one such comparator per output unit, the signed error is sufficient to train the connections it reaches directly, and the controls quantify the difficulties that appear when it is used, through fixed random feedback, to train earlier layers. Neither study shows that the motif is used this way in vivo. The unsigned control, moreover, establishes the need for directional information, not the superiority of this particular neuronal realisation over other ways of producing it.

In the language of loss functions, each comparator computes one term of the Hamming loss between input and reconstruction, *L*_*H*_ =Σ _*i*_ *x*_*i*_ *⊕ y*_*i*_, and F^+^ and F^*−*^ supply the polarity that the weight correction needs. The comparator is self-limiting: it is active while its own bit is wrong and silent once it is right. Every update is therefore corrective for its own unit, but this does not guarantee that the error of the whole network decreases monotonically, and in our networks reconstruction remains imperfect (18.4 wrong pixels per image on average). The decomposition into undershoot and overshoot maps naturally onto the separate populations of positive and negative prediction-error neurons reported in cortex (Keller & Mrsic-Flogel 2018, O’Toole et al. 2023), and it requires that, at the circuit level, the two components be routed to distinct plasticity pathways. The two feedback neurons of the signed-XOR motif provide distinguishable signals that could gate potentiation and depression; translating them into a change of synaptic strength, however, requires an additional plasticity rule, since an inhibitory neuron such as F^*−*^ does not depress a synapse merely by being inhibitory (Peña Fernández et al. 2026). Models of canonical interneuron circuits show how inhibitory plasticity can learn prediction-error neurons (Hertäg & Sprekeler 2020); the signed-XOR is a complementary, binary construction at single-neuron resolution, specified by its wiring rather than learned.

For the synapses it reaches directly, the signed comparator supplies exactly the error that a perceptron or delta rule requires (Rosenblatt 1962, Widrow & Hoff 1960), and this is sufficient to train a decoder that reconstructs its inputs. Each comparator considers only its own input and reconstruction, without knowledge of downstream consequences or of an overall cost function, yet the array of comparators produces accurate whole-image reconstructions. This supports the idea of distributed local learning, in which many small circuits, each performing signed XOR comparisons, collectively train a large system, and it resonates with classic error-correcting local rules (Rosenblatt 1962) and with Hebbian plasticity modulated by feedback signals (Stent 1973). What the present results add is the link between a specific microcircuit hypothesis and this computational function.

### 4.2 Credit assignment beyond the comparator

The control experiments delimit what the present model shows. The comparator solves the problem of computing a local error; it does not by itself solve the problem of assigning credit to synapses that are one layer removed from it. With fixed random sign feedback, encoder learning does not improve the reconstruction of digits when the hidden layer is at least as large as the input, and a fixed random encoder followed by a trained readout layer—the regime of random-feature and extreme learning machines (Huang et al. 2006)—performs better. The encoder rule does adapt the code to the trained digit classes when the hidden layer is narrow, although this adaptation does not transfer to EMNIST, and the gap to a gradient-trained network is moderate (about 4.6 balanced-accuracy points in the fully connected network and 8 in the convolutional one). This comparison is not matched in training budget or optimiser (3525 signed-XOR updates against 1175 Adam updates of a cross-entropy loss): it locates a gradient-trained network of the same architecture, not the isolated effect of using the gradient, and it supports no conclusion about relative efficiency. What could be read as a limitation of the model—that a fixed encoder works better—thus delimits experimentally the credit-assignment problem that the comparator leaves open. Closing it with a biologically plausible mechanism for hidden-layer credit assignment, for instance by letting comparators at intermediate stages compare top-down predictions with bottom-up activity, is the natural next step. These results also show why such claims need controls: reconstruction accuracy, transfer to EMNIST and linear decodability can all be obtained with a random encoder, and only comparisons against fixed-encoder, random-data and gradient-trained baselines reveal what the learning rule contributes.

### 4.3 Limitations and future directions

The model was deliberately simplified as a proof of feasibility: binary threshold neurons, discrete updates, a single hidden layer and small binary images. The autoencoders use the logical function of the comparators; the spiking dynamics of the signed-XOR motif have so far been studied only in isolated eight-neuron circuits (Peña Fernández et al. 2026), simulated in Brian2 (Stimberg et al. 2019), where its operation depends on the synaptic weights rather than requiring fast, parvalbumin-like inhibition. The next challenge is to embed spiking comparators in a learning network with a biophysical plasticity rule—for instance a three-factor rule gated by F^+^ and F^*−*^—that yields the same net effect as the bitwise rule used here.

Two roles of the feedback should also be distinguished (Peña Fernández et al. 2026). As a plasticity signal, the error projected onto the hidden layer changes the encoder weights and thereby the representations formed on later presentations; this is the role studied here, with a fixed random matrix. As a direct neural signal, F^+^ and F^*−*^ could also change the activity of the latent neurons they target without any immediate synaptic change. This dynamic feedback is a different mechanism and is not evaluated in this work.

The approach can also be extended to other learning paradigms. In supervised learning, an XOR motif could compare the network’s output with a teaching signal and drive learning, implementing a local delta rule; this requires a mechanism to supply the correct signal, for example feedback from higher areas or reward-related signals. This is consistent with evidence that burst-dependent plasticity can convey teaching signals from higher-level neurons and solve credit assignment in hierarchical circuits (Payeur et al. 2021). In sequence learning, XOR motifs could compare predicted and actual next sensory states, providing sequence prediction errors.

## 5 Conclusion

We have argued (Marco 2024) that identifying the biological analogues of network architecture, loss function and optimization algorithm would provide a unified account of learning in brains. This study tests the computational side of one such analogue for the loss function. The signed-XOR motif can produce a local error whose two components give each output unit the direction in which it must change (Peña Fernández et al. 2026); used as the only error signal, this signed error trains binary autoencoders without backpropagation through the synapses it reaches directly, and its sign is essential. Control experiments show that credit assignment to earlier layers with fixed random feedback remains limited. Together with the structural and spiking analysis of the motif, these results provide a network-level demonstration of feasibility—not a biological validation of the full process—for a concrete microcircuit as the source of the error, and a starting point for studying how such local errors could be propagated through deeper circuits.

## Declaration of competing interest

The authors have filed a European patent application relating to signed-XOR feedback for machine learning systems (European Patent Application No. 26382252.0).

## Data Availability Statement

MNIST and EMNIST are publicly available. All code used for the experiments, model fitting and plotting is available at https://github.com/MariaPFdez/XOR_classic and Zenodo Peña et al. (2026). The directory review_controls/ contains the control experiments: a re-implementation of the training rules that is bit-identical to the original code with default settings (verified by test_equivalence.py), the experiment grid, the per-run results for all configurations and seeds, and a script that prints every number reported in this article with the configuration it comes from.

## Funding

This work was supported by the Ministry for Digital Transformation and Public Administration, Spain [project CSIC-IASOMMA24-007], within the framework of the Recovery, Transformation and Resilience Plan and the Recovery and Resilience Facility, in accordance with Article 34(2) of Regulation (EU) 2021/241 of the European Parliament and of the Council. The funder had no role in study design, analysis, writing or the decision to submit.

## Acknowledgments

We thank the technical experts at the State Secretariat for Digitalization and Artificial Intelligence (Spain) for their support of this basic research topic.

The Instituto de Física de Cantabria (IFCA) has been accredited as Unidad de Excelencia María de Maeztu, a distinction awarded by the Spanish State Research Agency for the period 01/06/2026 - 31/05/2030.

## Notes

### Summary of Updates

The manuscript has been substantially revised to clarify the proposed XOR-based learning mechanism and its biological interpretation. The revised version introduces the signed-XOR formulation, providing directional error signals for synaptic updates, and clarifies its relationship to a complementary study investigating the neuronal circuit, its connectomic occurrence, and its spiking dynamics. The experimental analysis has been extended with additional validation across multiple random seeds and new control experiments, including comparisons with fixed random encoders, different network configurations, and gradient-based learning. These analyses allow a more precise interpretation of the results and clarify the specific contribution of the proposed mechanism. The discussion has also been revised to better distinguish local error computation from credit assignment in earlier network layers, explicitly acknowledge the limitations of the approach, and avoid overstating its biological implications. Finally, the methodology, figures, references, and reproducibility information have been updated and expanded.

